# Novel allelic variant of *Lpa1* gene associated with a significant reduction in seed phytic acid content in rice (*Oryza sativa* L.)

**DOI:** 10.1101/493981

**Authors:** D.S. Kishor, Choonseok Lee, Dongryung Lee, Jelli Venkatesh, Jeonghwan Seo, Joong Hyoun Chin, Zhuo Jin, Soon-Kwan Hong, Jin-Kwan Ham, Hee-Jong Koh

**Affiliations:** Department of Plant Science, Plant Genomics and Breeding Institute, and Research Institute of Agriculture and Life Science, Seoul National University, Seoul, South Korea; Graduate School of Integrated Bioindustry, Sejong University, Republic of Korea; Division of Biotechnology, Kangwon National University, Republic of Korea; Gangwon provincial Agricultural Research & Extension Services, Republic of Korea

## Abstract

In plants, *myo*-inositol-1,2,3,4,5,6-hexa*kis*phosphate (InsP_6_), also known as phytic acid (PA), is a major component of organic phosphorus (P), and accounts for up to 85% of the total P in seeds. In rice (*Oryza sativa* L.), PA mainly accumulates in rice bran, and chelates mineral cations, resulting in mineral deficiencies among brown rice consumers. Therefore, considerable efforts have been focused on the development of low PA (LPA) rice cultivars. In this study, we performed genetic and molecular analyses of *OsLpa*1, a major PA biosynthesis gene, in Sanggol, a low PA mutant variety developed via chemical mutagenesis of Ilpum rice cultivar. Genetic segregation and sequencing analyses revealed that a recessive allele, *lpa1-3*, at the *OsLpa*1 locus (Os02g0819400) was responsible for a significant reduction in seed PA content in Sanggol. The *lpa1-3* gene harboured a point mutation (C623T) in the fourth exon of the predicted coding region, resulting in threonine (Thr) to isoleucine (Ile) amino acid substitution at position 208 (Thr208Ile). Three-dimensional analysis of Lpa1 protein structure indicated that *myo*-inositol 3-monophosphate [Ins(3)P_1_] kinase binds to the active site of Lpa1, with ATP as a cofactor for catalysis. Furthermore, the presence of Thr208 in the loop adjacent to the entry site of the binding pocket suggests that Thr208Ile substitution is involved in regulating enzyme activity via phosphorylation. Therefore, we propose that Thr208Ile substitution in *lpa1-3* reduces Lpa1 enzyme activity in Sanggol, resulting in reduced PA biosynthesis.

## Introduction

In most cereal crops, *myo*-inositol-1,2,3,4,5,6-hexa*kis*phosphate (InsP_6_), also known as phytic acid (PA), is considered a major source of phosphorus (P) available in the form of phytate, and accounts for 65–85% of the total P in seeds [1]. Monogastric animals poorly digest PA, as they lack the phytase enzyme, which is responsible for the release of phosphate residues [2]. PA is an efficient chelator of mineral cations, such as zinc (Zn^2+^), iron (Fe^2+^), magnesium (Mg^2+^), potassium (K^2+^), and calcium (Ca^2+^), in the nutritional tract. Because of these attributes, PA is considered as an antinutrient [3, 4]. Hence, there is a need to develop low PA (LPA) crop cultivars to maximize the nutritional benefits of grains.

Mutants associated with the LPA phenotype have been identified in several crop plants including maize (*Zea mays*) [5, 6], barley (*Hordeum vulgare*) [7], soyabean (*Glycine max*) [8], rice (*Oryza sativa*) [9], and wheat (*Triticum aestivum*) [10]. Although, LPA mutants are identified primarily on the basis of percentage reduction of PA and high inorganic P (P_i_) content in seeds [5, 11], some mutants show a significant accumulation of *myo*-inositol and inositol phosphate [Ins(1,3,4)P_3_ 5-/6] intermediates in seeds [12, 13].

Previously, the LPA phenotype of seeds has been associated with reduced agronomic performance of mutant crop plants in the field [5, 14]. It is important to understand the genetic and molecular bases of reduced agronomic performance of LPA mutants for effective utilization in breeding programs. In addition, studies show that climate change and elevated carbon dioxide (CO_2_) levels negatively affect micronutrient bioavailability and total P in grains [15, 16]. Therefore, developing crop cultivars with increased micronutrient bioavailability in seeds and greater adaptability to environmental variations, by reducing the PA content in grains, is an important priority of breeding programs.

PA is biosynthesized via two different routes: lipid dependent and lipid independent [3, 17]. The lipid dependent pathway operates in all plant organs, whereas the lipid independent pathway is predominant only in seeds [13, 17, 18]. In the first step of PA biosynthesis, D-glucose-6-phosphate is converted to *myo*-inositol 3-monophosphate [Ins(3)P_1_] by *myo*-inositol 3-phosphate synthase (MIPS) [19]. This is followed by the sequential phosphorylation of specific inositol to InsP_6_ through the action of various inositol phosphate kinases (S1 Fig). However, enzymes involved in lipid independent PA biosynthesis, from Ins(3)P_1_ seem to be complicated and are not well understood [3]. Nevertheless, PA biosynthetic genes encoding other *myo*-inositol enzyme and inositol phosphate kinases are well documented in major plants [12, 13, 20, 21]. Additionally, biochemical and functional analyses of PA biosynthetic genes encoding Ins monophosphate kinase could address the missing steps in the lipid independent pathway.

In rice, several mutants with low seed PA content have been reported [14, 21–27]. Genetic studies of LPA mutants have shown that a single recessive gene is responsible for the LPA phenotype in rice and other crop plants [21, 22, 27, 28]. The first *lpa* gene encoding inositol 1,3,4-triskisphophate 5/6-kinase (ITPK5/6) was identified in maize, and designated as *Lpa2*. Subsequently, *myo*-inositol kinase gene *Lpa3*, and multidrug resistance protein (MRP) ATP binding cassette (ABC) transporter gene *Lpa1* were identified [12, 13, 29]. In addition, reduction of PA content in *Arabidopsis atipk2β* mutant indicates the inositol 1,4,5-tris-phosphate (IPK2) kinase of lipid dependent pathway is also active the seeds [20]. In rice, *OsLpa*1 gene has been associated with the reduction in seed PA content and increase in seed P_i_ content, with little change in the total P content in seeds [22, 30]. *OsLpa*1 have homology with one gene, Os09g0572200 (*OsLpa*1 paralog) within the rice genome, suggests possible overlapping or redundant functions [22].

Genetic studies in rice have confirmed that a mutation in the *OsLpa*1 locus generates the LPA phenotype in seeds. Molecular characterization of LPA mutants has previously revealed three alleles of the *OsLpa*1 locus, including KBNT *lpa 1-1*, DR1331-2, and Os-lpa-XQZ-1, responsible for the low PA phenotype of seeds [22, 30]. In the present study, we report a novel allele of *OsLpa*1, *OsLpa1-3*, responsible for a significant reduction in the seed PA content in a new LPA mutant rice cultivar Sanggol developed in the Republic of Korea [31]. Sequence analysis of *OsLpa*1-3 revealed a point mutation in the gene coding sequence. Our data suggest that this mutation is responsible for the LPA phenotype of Sanggol mutant.

## Material and methods

### Plant material

The low PA mutant rice cultivar Sanggol derived from a *japonica* rice cultivar Ilpum mutagenized with *N*-methyl-*N*-nitrosourea (MNU) [31]. Ilpum was used as the wild type in comparing phenotypic data. Sanggol was crossed with Ilpum to develop F_2_ population. Segregation analysis was performed using the F_2_ population. Both parent cultivars and F_2_ individuals were grown in experimental fields of Seoul National University, Republic of Korea.

### Agronomic trait analysis

To characterize agronomic traits, 15 phenotypic observations were recorded during various stages of plant growth, according to the Standard Evaluation System (SES) for rice, 2014. Yield data was obtained from “3.6 m X 3.6 m” plot size. All agronomic data were analyzed using the Student’s *t*-test in SPSS 16.0 (https://www.ibm.com/analytics/spss-statistics-software) to determine significant differences among Sanggol, and Ilpum cultivars.

### Analysis of P_i_ and PA content in seeds

Concentrations of P_i_ and PA in seeds were examined using P^31^ nuclear magnetic resonance (P^31^ NMR) spectroscopy [32], with slight modifications.

#### Sample preparation

Fine powdered samples (1 g dry weight) of brown rice were thoroughly mixed with 10 mL of 2.4% HCl in 14 mL Falcon tubes. Samples were incubated at room temperature for 16 h on an HB-201SF shaker (HANBAEK Scientific Co) at 220 rpm, and subsequently centrifuged at 1,500 × *g* (combi 514R, Hanil science Inc.) at 10°C for 20 min. Crude extracts were transferred to a new 14 mL Falcon tube containing 1 g NaCl, and incubated at 25°C for 40 min on a shaker at 220 rpm to dissolve NaCl. Samples were allowed to settle at 4°C for 60 min, and then centrifuged at 1,500 × *g* at 10°C for 20 min.

#### ^31^P NMR

For ^31^P NMR spectroscopy, samples were prepared by mixing 450 μL of NaCl treated acid extract with 450 μl of buffer containing 0.11mM EDTA-disodium salt and 0.75 mM NaOH, 40 mg NaOH, and 100 μL D_2_O in 1.5 mL microtubes. Sample and standard peaks were obtained on a 600 MHz spectrometer using Advance 600 ^31^P NMR system (Bruker, Germany). PA sodium salt and 85% phosphoric acid were used as external standards for peak identification and further analysis [33, 34]. For internal calibration, 1 mM of phenylphosphonic acid was included in 100 μL D_2_O during NMR measurements. All standards were purchased from Sigma-Aldrich, USA.

To determine significant differences in seed PA and P_i_ contents among parents and F_2_ individuals, data were analyzed using the Student’s *t*-test in SPSS 16.0 (https://www.ibm.com/analytics/spss-statistics-software).

### Expression analysis of PA biosynthetic genes

Genes involved in PA biosynthesis and transport were identified from the RAB-DB and from recent studies [25, 35, 36]. The rice microarray database, RiceX-Pro, shows different expression patterns of most of the PA biosynthetic genes in various tissues and organs [37]. To confirm the expression pattern of PA biosynthetic genes, spikelets were harvested from the wild cultivar Ilpum at 5 days after flowering (DAF), and total RNA was extracted using RNAiso Plus (Takara Bio, Japan). The extracted RNA samples were treated with RNase-free recombinant DNase Ι (Takara Bio, Japan) to eliminate genomic DNA contamination, and first-strand cDNA was synthesized using M-MLV reverse transcriptase (Promega, USA). The PA biosynthetic genes (200–550bp) were amplified from cDNA samples by reverse transcription polymerase chain reaction (RT-PCR) using gene-specific primers (Table 1) with the following conditions: initial denaturation at 95°C for 2 min, followed by 32 cycles of denaturation at 95°C for 20 s, annealing at 58°C for 40 s, and extension at 72°C for 1 min, and a final extension at 72°C for 5 min. The *Actin* gene was used as an internal control.

**Table 1.**
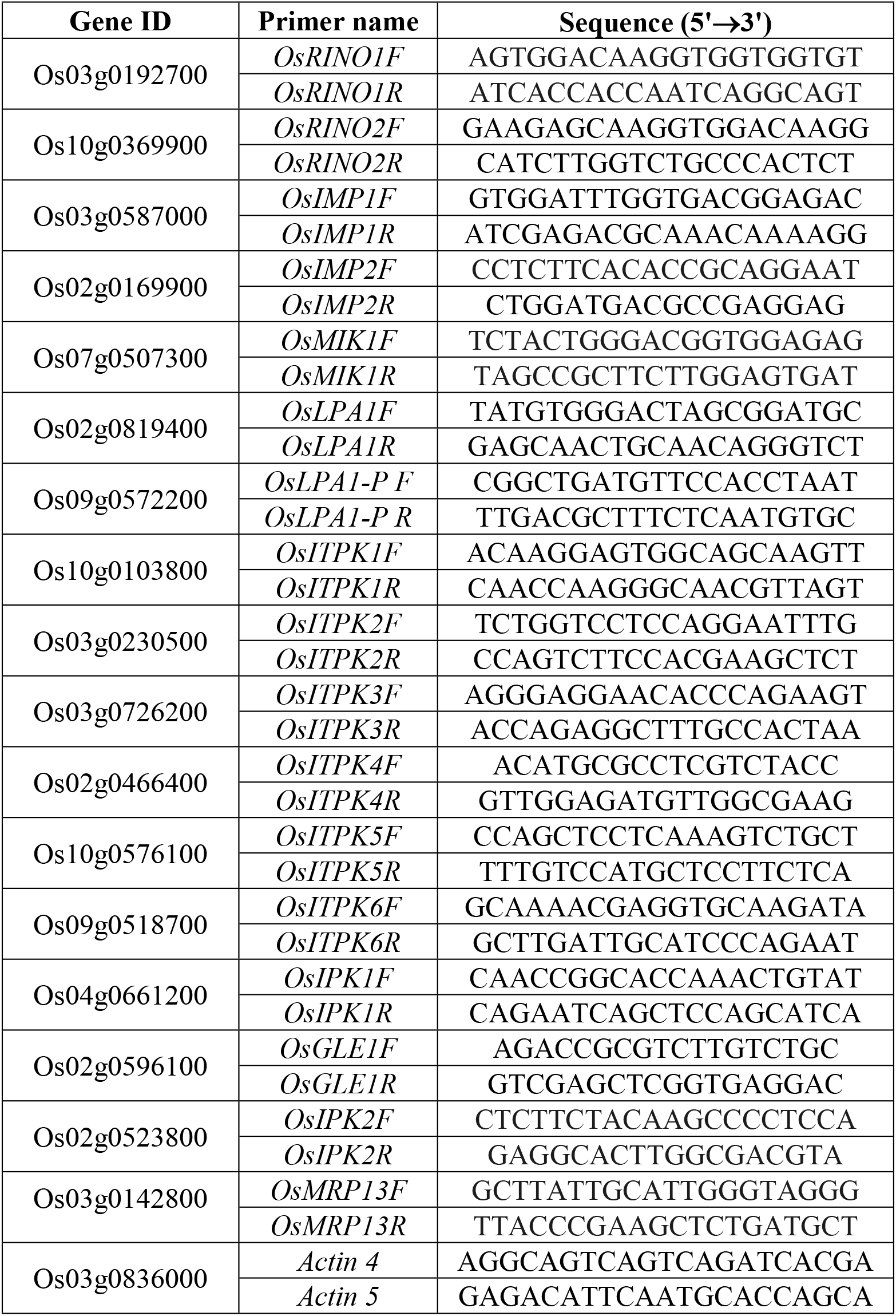
RT-PCR primers used to amplify PA biosynthetic genes.

### Sequence analysis

Genomic DNA and cDNA were isolated from young leaves and spikelets of Sanggol low PA mutant cultivar, respectively. Fragments of size 300bp-2000bp were amplified from the coding region and untranslated region (UTR) of 16 genes of lipid dependent and independent pathways using gene-specific primers designed with prime3 (http://bioinfo.ut.ee/primer3-0.4.0/) (S1 Table). The PCR products were purified using the DNA Purification Kit (Inclone, Korea), and analyzed with an ABI Prism 3730 XL DNA Analyzer (PE Applied Biosystems, USA). In addition, sequences of all 16 genes in Ilpum was downloaded from the crop molecular breeding lab server (http://nature.snu.ac.kr/rice/). Sequences were aligned using the Codon Code Aligner software (Codon Code Corporation, USA).

### Candidate gene analysis

To confirm nucleotide polymorphisms in the candidate genes, validation primers were designed using Primer3 for cDNA sequencing (Table 2). The PCR products were purified using the DNA Purification Kit (Inclone, Korea), and analyzed with an ABI Prism 3730 XL DNA Analyzer (PE Applied Biosystems, USA). Sequences were aligned using the Codon Code Aligner software (Codon Code Corporation, USA). Simultaneously, BLAST search was performed using the predicted amino acid sequences of the candidate genes in the NCBI database (https://blast.ncbi.nlm.nih.gov/Blast.cgi), and deleterious amino acid substitutions were predicted using Provean web server with proven scores [38].

**Table 2.**
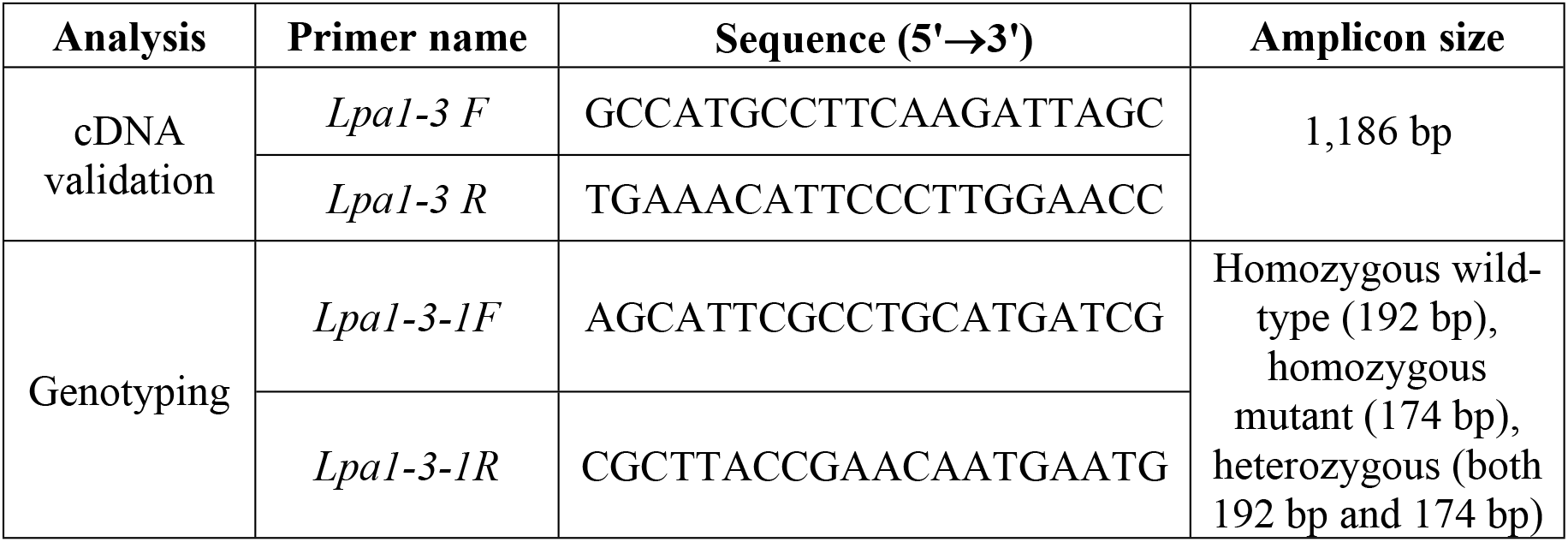
Primers used for validating cDNA sequences and genotyping the F_2_ population.

### Expression analysis of *Lpa* and lipid dependent PA biosynthesis genes in Sanggol and Ilpum cultivars

Total RNA was extracted from the leaves at 15 days after germination (DAG) to analyze the expression of *Lpa* and lipid dependent pathway genes, and 5 DAF from spikelets to analyze the expression of *OsLpa*1 in Sanggol and Ilpum cultivars. For the expression analysis of *OsLpa*1 paralog and *OsIpk*2 genes, total RNA was extracted only from spikelets at 5 DAF. RNA extraction was performed as described above. The extracted RNA was subjected to RT-PCR using gene-specific primers (Table 3). The *Actin* gene was used as an internal control.

**Table 3.**
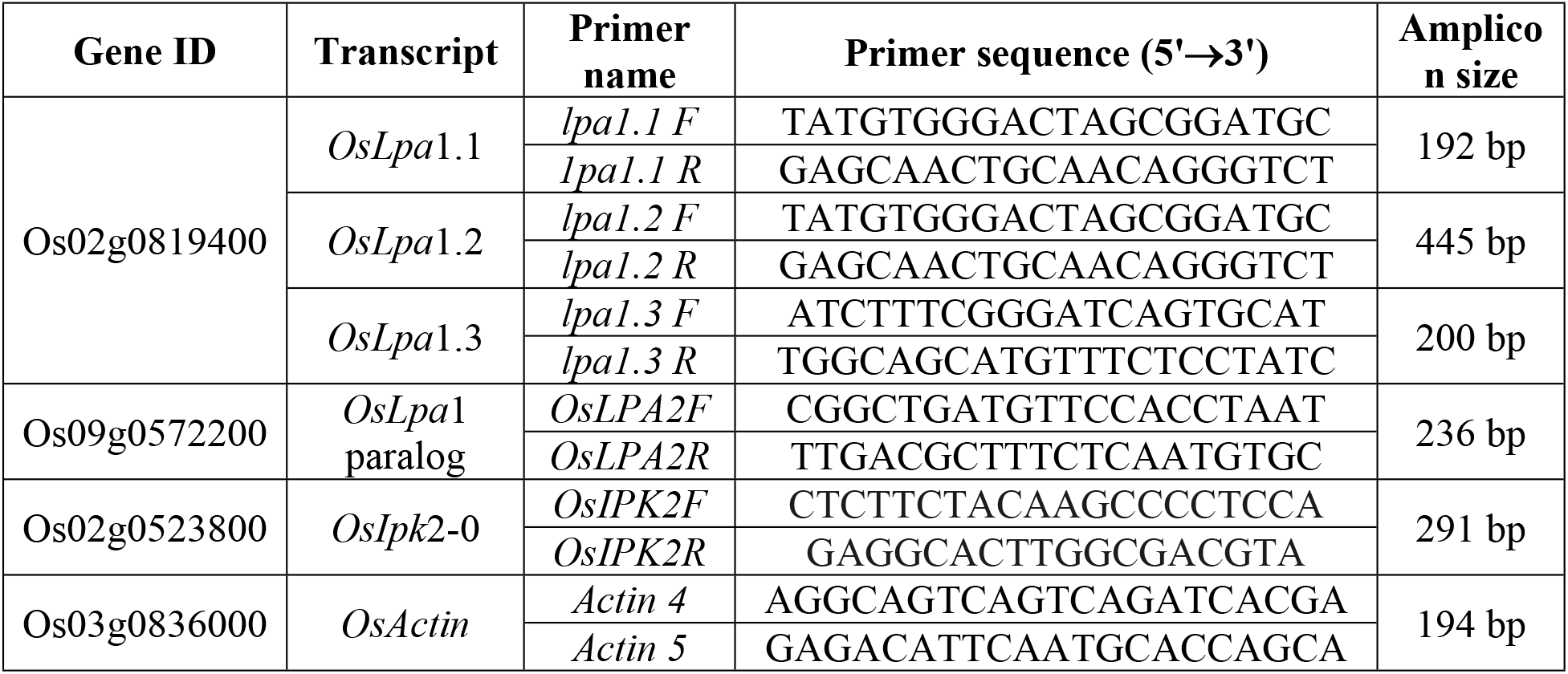
RT-PCR primers used to amplify *Lpa* and *Ipk2* genes.

### Derived cleaved amplified polymorphic sequence (dCAPS) analysis

Genomic DNAs were isolated from all 96 F_2_ plants derived from cross between Sanggol and Ilpum, and subjected to dCAPS analysis. The F_2_ genotyping primers (Table 2) were designed using dCAPS 2.0 (http://helix.wustl.edu/dcaps/) to validate a single nucleotide substitution (C to T) in the *OsLpa*1 gene in Sanggol cultivar, which generates a *Taq*I restriction site (TCGA) in the amplified PCR product. PCR was performed using the following conditions: initial denaturation at 95°C for 2 min, followed by 32 cycles of denaturation at 95°C for 20 s, annealing at 58°C for 40 s, and extension at 72°C for 30 s, and a final extension at 72°C for 1 min. The amplified PCR product was digested with *Taq*I restriction endonuclease (Promega, USA), and separated on 3% agarose gel.

### Multiple sequence alignment and phylogenetic analysis

Amino acid sequences of the Lpa superfamily were obtained from the NCBI protein database (https://blast.ncbi.nlm.nih.gov/Blast.cgi?PAGE=Proteins), and subjected to multiple sequence alignment using the Clustal Omega program (https://www.ebi.ac.uk/Tools/msa/clustalo/). Multiple sequence alignment editing, visualization, and analysis was performed using Jalview 2.10.4 (http://www.jalview.org/). The Lpa and other superfamily proteins obtained from the NCBI protein database were used for phylogenetic analysis. Neighbour-joining tree was constructed using MEGA 7 [39] with 1,000 bootstrap replicates.

### Biocomputational analysis

A three-dimensional (3D) model of Lpa1 protein was produced under the intensive mode of the Phyre2 server [40] (www.sbg.bio.ic.ac.uk/phyre2/html/). The ligand and cofactor were downloaded from the PubChem database (https://pubchem.ncbi.nlm.nih.gov/) for protein ligand analysis. Furthermore, auto docking and 3D model were analyzed using the CLC drug discovery workbench 4.0 (QIAGEN, Denmark). Putative phosphorylation sites were predicted with the GPS 3.0 server (http://gps.biocuckoo.org/) using high cut-off values ranging from 1.36 to 17.72.

## Results

### Agronomic characterization of Sanggol low PA mutant and Ilpum cultivars

Analysis of agronomic traits demonstrated a significant reduction in the plant height (cm), number of productive tillers, culm length (cm), first intermodal length (cm), 1,000-grain weight (g), number of spikelets per panicle, number of panicles per plant, and yield components of Sanggol compared with Ilpum (Table 4 and Fig 1A). By contrast, the number of days to 50% flowering was significantly higher in Sanggol than in the Ilpum, indicating delayed flowering in the mutant cultivar. In addition, Sanggol exhibited significantly higher percentage of chalky grains compared with the wild cultivar. However, no significant differences were observed between the two cultivars in morphological characteristics, such as secondary internodal length, grain length, grain width, panicle length, and spikelet fertility (Fig 1B and Fig 1C). Overall, these data indicate that Sanggol low PA mutant shows poor agronomic performance with respect to the flowering time, yield, and yield components compared with the Ilpum.

**Table 4.** Agronomic traits of the wild-type cultivar Ilpum and low PA mutant cultivar Sanggol.

| **Traits<sup>a</sup>** | **PH**<br>**(cm)** | **NPT** | **DFF** | **CL**<br>**(cm)** | **FIL**<br>**(cm)** | **SIL**<br>**(cm)** | **GL**<br>**(mm)** | **GW**<br>**(mm)** | **KGW**<br>**(g)** | **GC**<br>**(%)** | **NSP** | **NPP** | **PL**<br>**(cm)** | **SPF**<br>**(%)** | **YPP**<br>**(g)** |
| --- | --- | --- | --- | --- | --- | --- | --- | --- | --- | --- | --- | --- | --- | --- | --- |
| Wild | 97 | 22 | 116.7 | 75 | 40.5 | 15 | 5.01 | 2.97 | 20.93 | 43.62 | 187 | 12 | 22 | 97 | 760 |
| Mutant | 77 | 13 | 122.5 | 54 | 35.5 | 14.83 | 4.81 | 3.03 | 18.9 | 54.66 | 165 | 8 | 23 | 96 | 470 |
| Significance<sup>b</sup> | ** | ** | ** | ** | ** | NS | NS | NS | ** | ** | ** | * | NS | NS | ** |

<sup>a</sup>Traits: PH, plant height; NPT, number of productive tillers; DFF, days to 50% flowering; CL, culm length; FIL, first internodal length; SIL,
second internodal length; GL, grain length; GW, grain width; KGW, 1,000 grain weight; GC, grain chalkiness; NSP, number of spikelets per
panicle; NPP, number of panicles per plant; PL, panicle length; SPF, spikelet fertility; YPP, yield per plot
<sup>b</sup>Asterisks indicate the level of significance (\*, $P < 0.05$ ; \*\*, $P < 0.01$ ). NS, non-significant.

**Fig 1.**
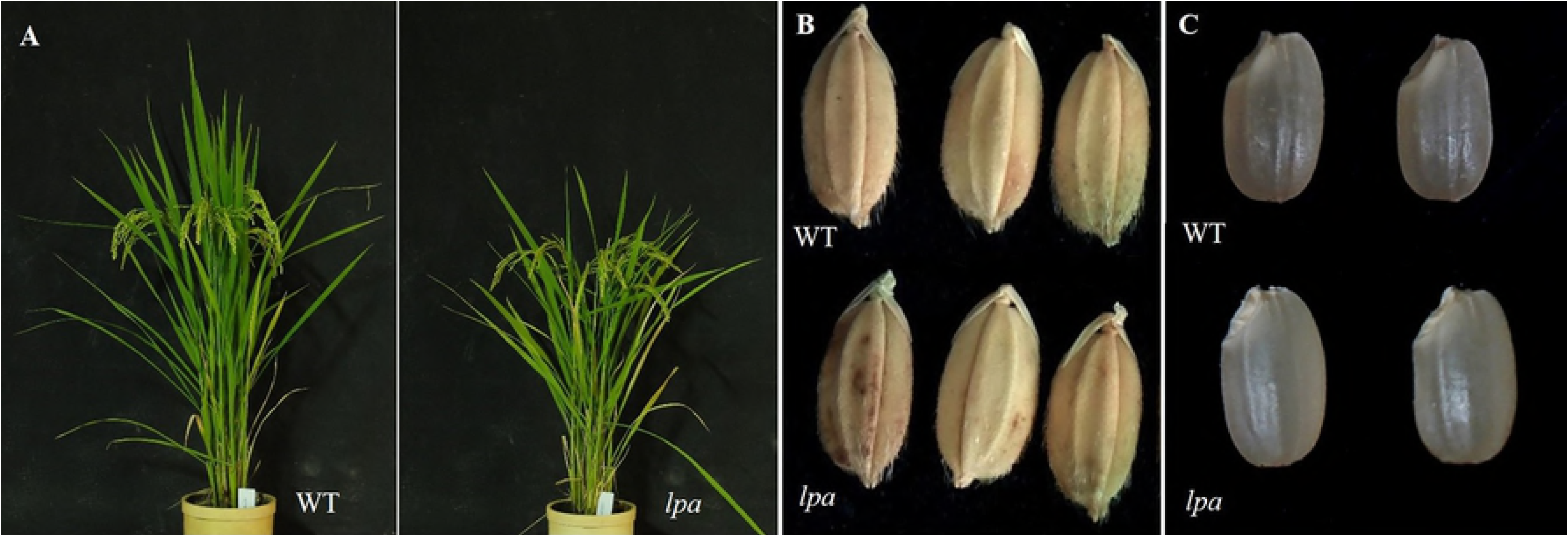
Phenotypic comparison between Sanggol mutant (*lpa*) and wild-type (WT) cultivar Ilpum. (A) Whole plant. (B) Spikelet. (C) Mature grain.

### Determination of PA and P_i_ content in Sanggol and Ilpum seeds

To quantify PA and P_i_ content in seeds, brown rice extracts of Sanggol and Ilpum were analyzed via ^31^P NMR spectroscopy. Results showed that PA contents were significantly reduced (49% reduction), and P_i_ content was significantly increased in the seeds of Sanggol compared with Ilpum (Table 5). The ^31^P NMR analysis showed peaks analogous to standard (Fig 2A) for P_i_ and PA peak identification. Similarly, Pi and PA analogous peaks were observed for wild (WT) (Fig 2B), and mutant (*lpa*) (Fig 2C) types.

**Table 5.** Seed PA and P_i_ content in Sanggol and Ilpum cultivars.

| Cultivar | PA P (mg g<sup>-1</sup>) | P<sub>i</sub> (mg g<sup>-1</sup>) | Total P (mg g<sup>-1</sup>) |
| --- | --- | --- | --- |
| Ilpum | 5.7 ± 0.34 | 0.1 ± 0.04 | 5.85 ± 0.34 |
| Sanggol | 2.9 ± 0.69* | 1.8 ± 0.10** | 5.21 ± 0.62 |

**Fig. 2.**
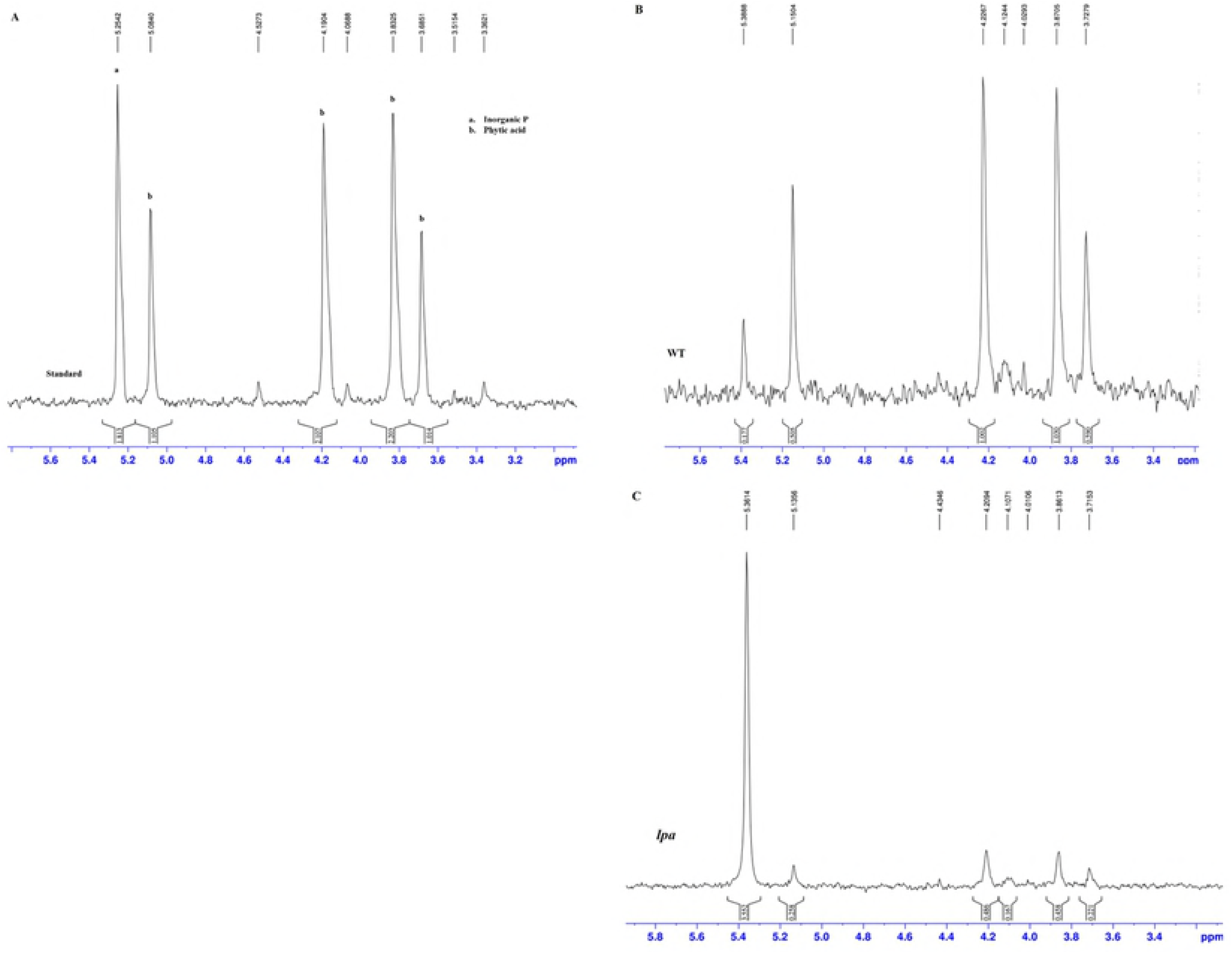
^31^P NMR spectrum of standard, Ilpum (WT), and Sanggol mutant (*lpa*) (A) Reference standard peaks. (B) Wild ‘WT’ (C) *lpa* ‘Mutant’.

Additionally, PA and P_i_ amounts were also quantified among 96 F_2_ individuals using ^31^P NMR spectroscopy. Segregation analysis revealed that 77 F_2_ plants showed the wild-type phenotype, whereas 19 F_2_ plants showed the mutant phenotype (Table 6), and the phenotype segregation fitted a 3:1 ratio, suggesting that a single recessive allele controls the low PA in the seeds of Sanggol mutant cultivar.

**Table 6.**
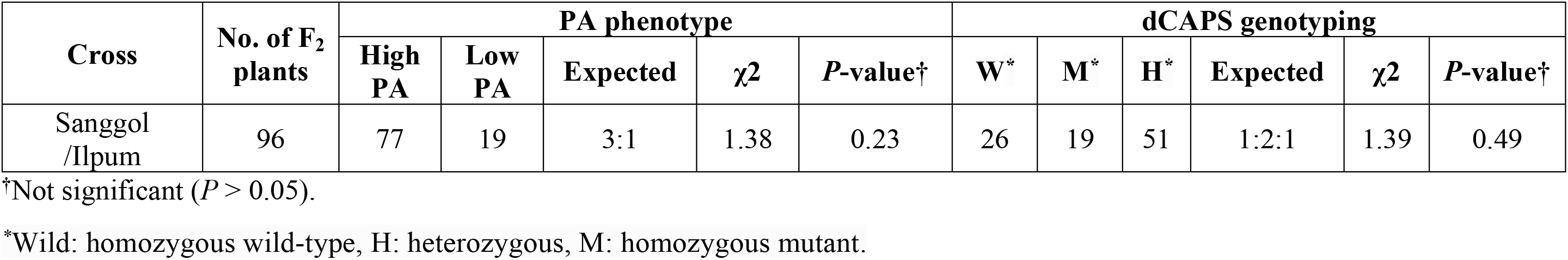
Segregation and co-segregation analysis of seed PA content among 96 F_2_ individuals derived from a cross between Sanggol and Ilpum cultivars.

### Expression of PA biosynthetic gene and sequence analysis

To identify the gene responsible for reduced PA content in seeds, the candidate gene approach was followed. In rice, PA biosynthesis and accumulation begins after flowering [42, 43], and continues until 25 DAF during seed development [44]. Therefore, we extracted total RNA from ‘Ilpum’ spikelets at 5 DAF, and subjected it to RT-PCR analysis. Results showed that 15 genes in the PA biosynthesis pathway were expressed at 5 DAF (Fig 3). Further, we amplified and sequenced 16 genes involved in PA biosynthesis from sanggol and Ilpum cultivars (S1 Table).

**Fig 3.**
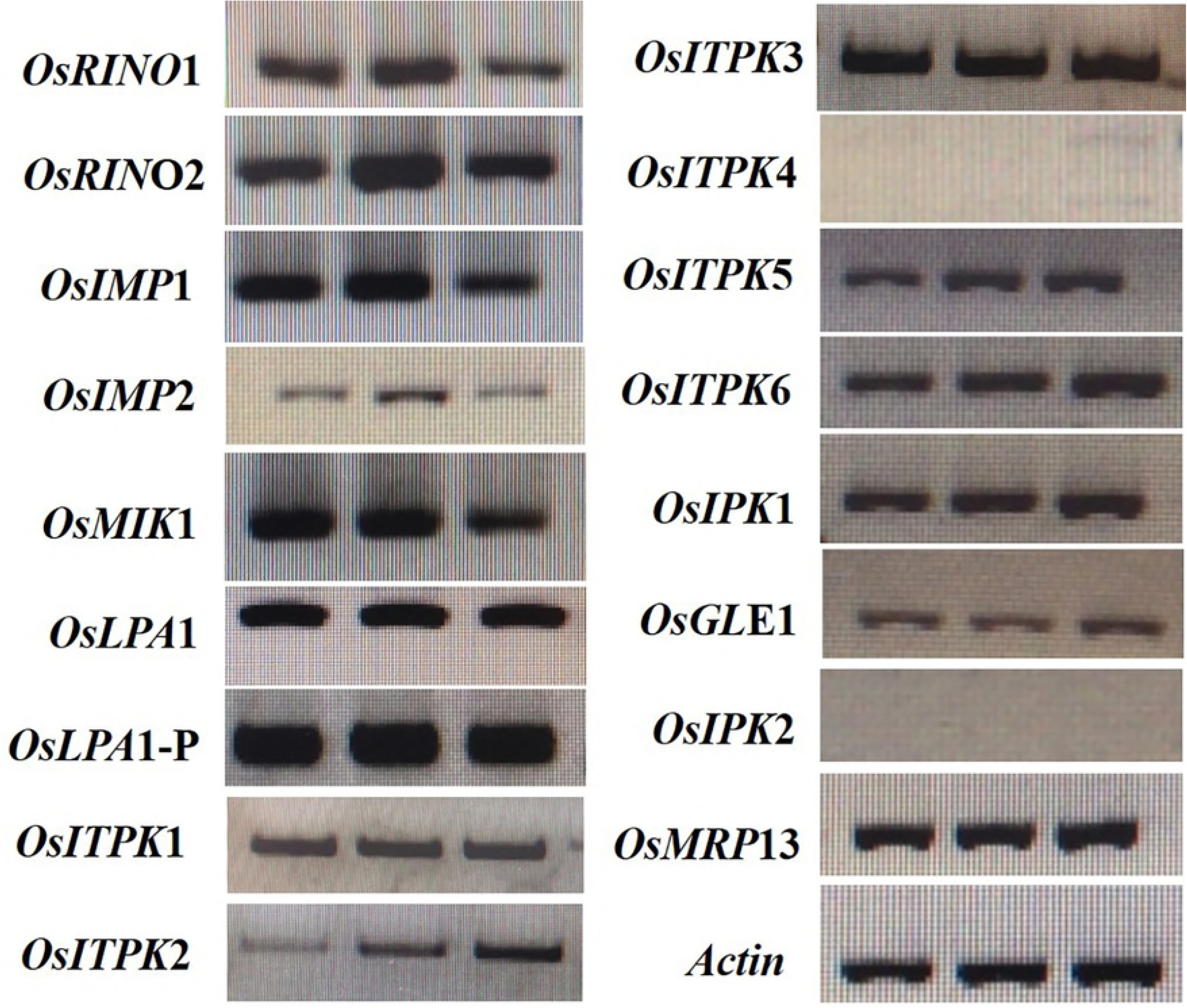
Semi-quantitative RT-PCR analysis of PA biosynthetic genes at 5 DAF in the Ilpum cultivar.

Sequence analysis of PA biosynthetic genes revealed a single nucleotide polymorphism (SNP) in the *OsLpa*1 gene of Sanggol *lpa* mutant (Fig 4A); none of the other PA biosynthetic genes showed mutations in Sanggol *lpa* mutant. Previously, the *OsLpa*1 locus has been mapped to chromosome 2 [11], and narrowed down to a region less than 150 kb using microsatellite and sequence tagged site markers [45]. Further, the *OsLpa*1 has been characterized in *lpa* mutants of rice [22, 30]. The *OsLpa*1 gene encodes three expressed splice variants in rice [22, 35]. Sequence analysis of the *OsLpa*1 locus (position +1 to 2,058 bp; Genbank accession number: MH707666) showed a SNP (C623T) in the fourth exon of the largest splice variant, designated as *OsLpa*1-3.1, in Sanggol *lpa* mutant. Additionally, another SNPs (C53T) was identified in the first exon of the small splice variants, *OsLpa*1-3.2 and *OsLpa*1-3.3 (S2 Fig). Further, sequence analysis of *OsLpa*1-3.1 cDNA confirmed the presence of *lpa1*-3 allele in Sanggol mutant (Fig 4B).

**Fig 4.**
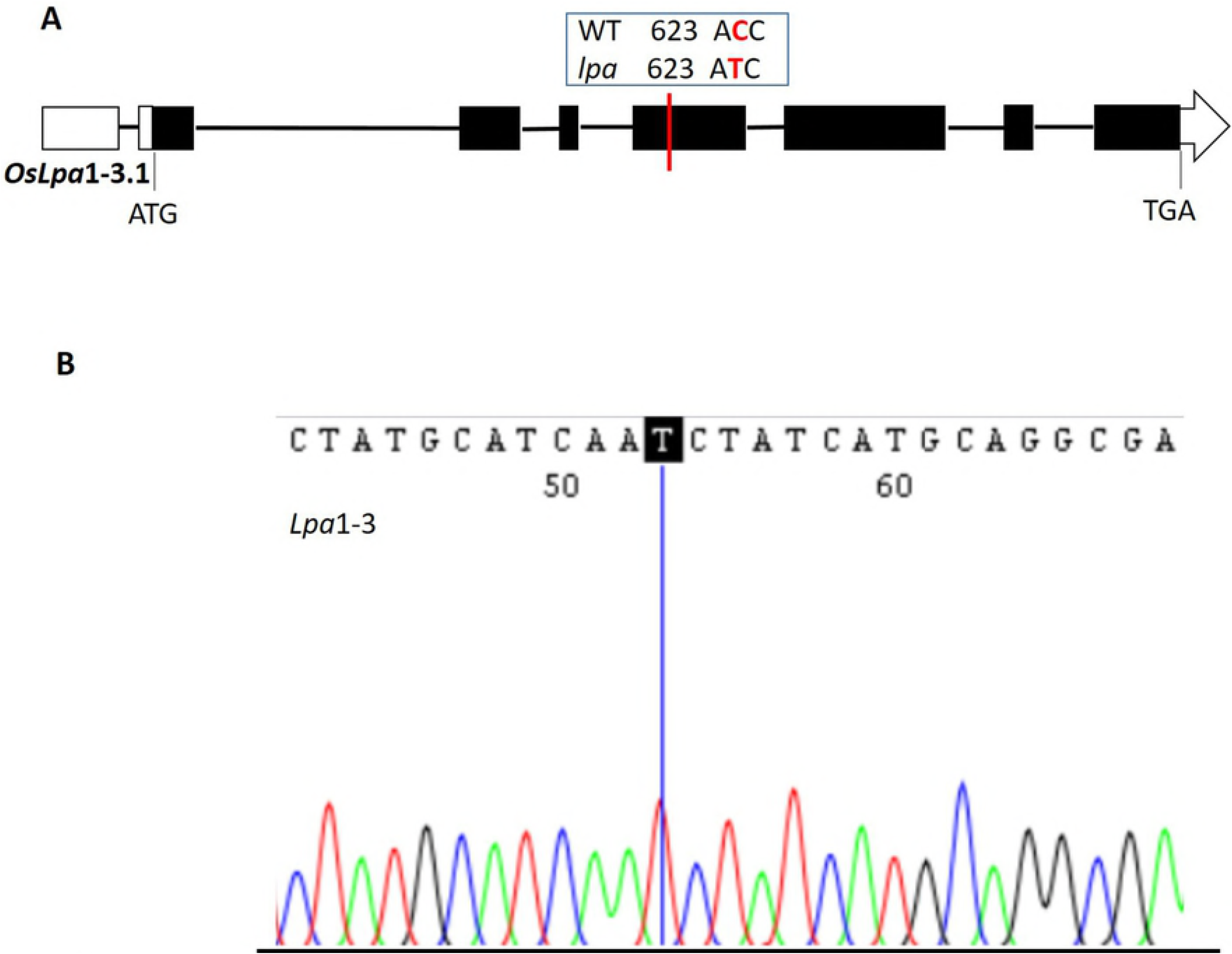
Gene structure of *OsLpa*1 according to Kim et al. [22] and Zhao et al. [30]. (A) Structure of the novel splice variant *OsLpa*1-3.1 carrying the C623T mutation in Sanggol low PA mutant cultivar. Empty boxes represent 5’UTR and 3’UTR, black box represents the coding region, and lines between boxes indicate introns. ATG (start codon) and TGA (stop codon) are shown for each splicing variant. (B) cDNA validation of *OsLpa*1-3.1 showing *Lpa*1-3 allele in the Sanggol mutant cultivar.

To determine the expression of *OsLpa*1 splice variants in mutant and wild-type cultivars, we performed RT-PCR analysis of *OsLpa*1 gene at 15 DAG using total RNA isolated from leaves and spikelets at 5 DAF. Expression analysis revealed that both *OsLpa*1-3.1 and *OsLpa*1-3.2 were expressed at 15 DAG, with slightly different expression patterns, whereas *OsLpa*1-3.3 showed no expression at 15 DAG in both cultivars (Fig 5A), indicating that *OsLpa*1-3.1 and *OsLpa*1-3.2 play an important role in seedling growth. At 5 DAF, *OsLpa*1-3.1 showed strong expression in both Sanggol *lpa* mutant and wild cultivar Ilpum; however, *OsLpa*1-3.3 exhibited low expression in both cultivars, and *OsLpa*1-3.2 exhibited no expression in either cultivar, suggesting *OsLpa*1-3.1 as a candidate transcript responsible for the seed low PA phenotype of Sanggol mutant. Protein analysis of Lpa1 amino acid sequence predicted deleterious amino acid substitution changes threonine (Thr) to isoleucine (Ile) in *OsLpa*3.1 (Thr208Ile), with a -5.715 proven score. Similarly, deleterious amino acid substitution changes were observed in *OsLpa*3.2 and *OsLpa*3.3 (Thr18Ile), with -5.482 proven scores.

**Fig 5.**
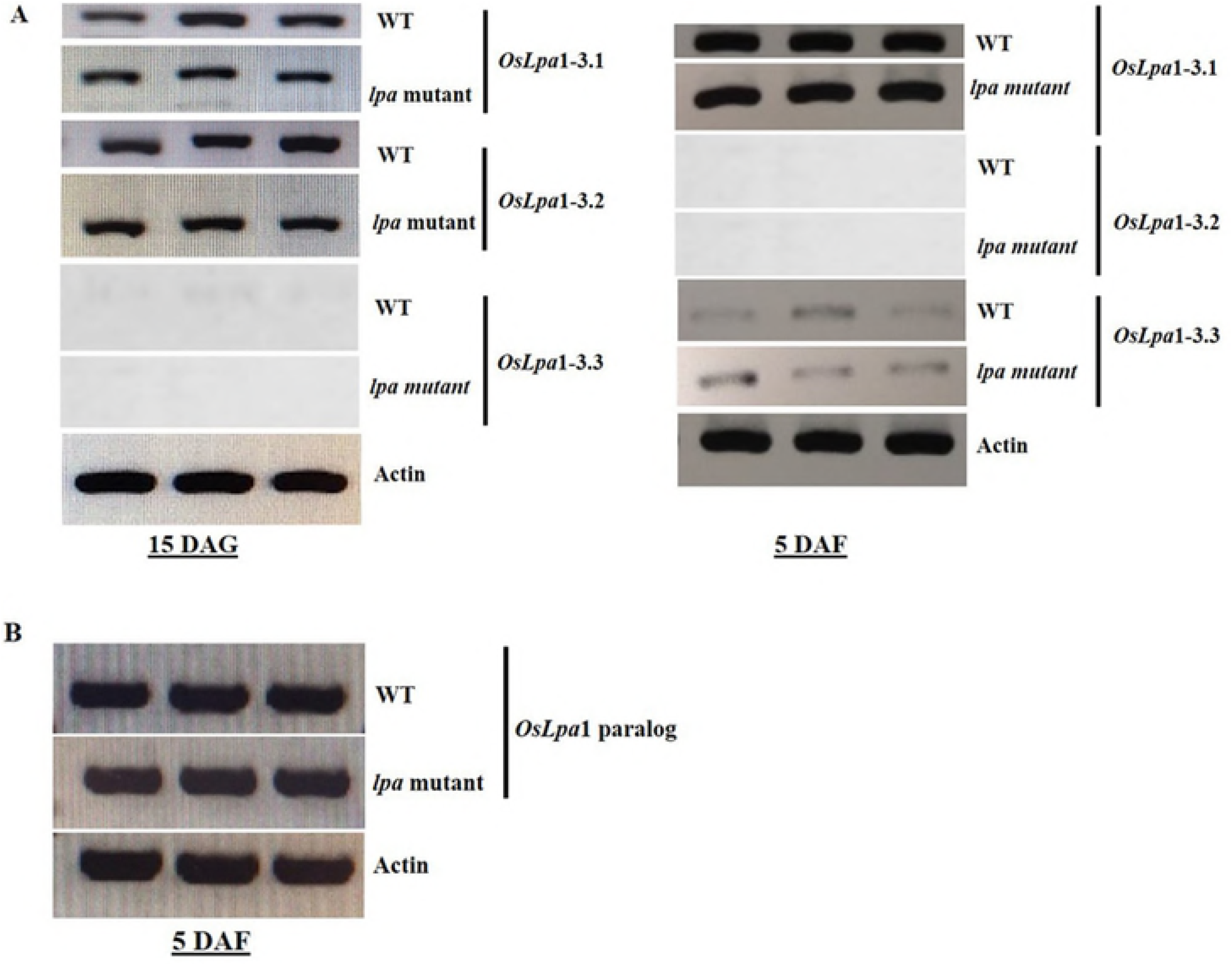
RT-PCR analysis of the *Lpa* gene family in Sanggol and Ilpum. (A) Expression of *OsLpa*1 gene at 15 DAG and 5 DAF. (B) Expression of *OsLpa*1 gene paralog at 5 DAF.

Additionally, expression of the *OsLpa*1 paralog, reported previously by Kim et al. [22], was investigated at 5 DAF in Sanggol *lpa* mutant and wild cultivars using RT-PCR. The *OsLpa*1 paralog exhibited strong expression in both Sanggol *lpa* mutant and wild cultivars (Fig 5B), suggesting that sequence variation in the coding region of *OsLpa1* was responsible for the low PA content of Sanggol seeds. In addition, reduction of PA content in *Arabidopsis atipk2β* mutant indicates the IPK2 kinase of lipid dependent pathway is active the seeds [20]. We also ruled out the possibility for seed PA biosynthesis similar to *Arabidopsis* in Sanggol low PA mutant cultivar. However, our RT-PCR results showed no expression of *OsIpk2*, a key PA biosynthesis gene in the lipid dependent pathway (data not shown), suggesting that the lipid dependent pathway is not active in the Sanggol or Ilpum cultivar.

Next, we performed multiple sequence alignment of Lpa1 amino acid sequences of Sanggol and other major plant species. Results revealed an amino acid substitution in the conserved kinase domain in Sanggol (Fig 6A), thus showing the impact of a SNP in gene coding sequence. The kinase domain of Lpa1 shows weak homology with that of 2-phosphoglycerate kinase (2-PGK) found in hyperthermophilic methanogens [22]. However, there is structural similarity among the substrates and products of 2-PGK and Lpa1 [46].

**Fig 6.**
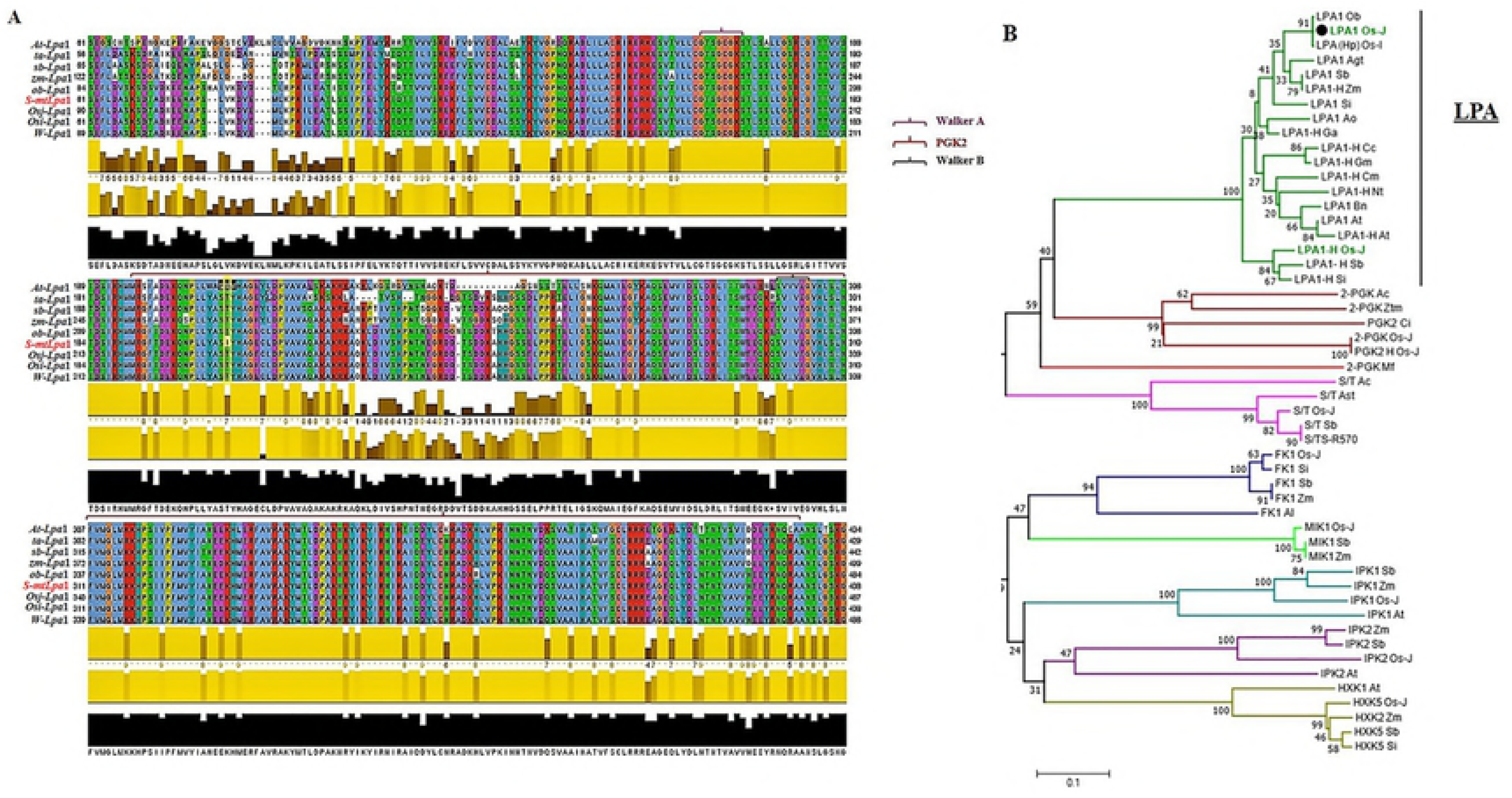
Multiple sequence alignment and phylogenetic analysis of Lpa protein. (A) Multiple sequence alignment of Lpa1 protein from Sanggol (s-mtLpa1) and other major plant species. The P-loop kinase domain of 2-PGK found in *Methanothermus fervidus* is marked with a red line. P-loop is an ATP/GTP binding site motifs (Walker A and Walker B) are indicated using pink and black arrowheads, respectively. The yellow box shows a single amino acid substitution in the conserved kinase domain in Sanggol cultivar. (B) Phylogenetic relationship among protein families of various kinases, including 2-PGK, serine threonine protein kinase (S/T), fructokinase (FK), *myo*- inositol kinase 1 (MIK), inositol 1,3,4,5,6 penta*kis*phosphate 2-kinase (IPK1), inositol 1,4,5-tris-phosphate kinase (IPK2), hexokinase 1 (HXK), and hexokinase 2, with the Lpa protein clade. Phylogenetic tree was constructed using amino acid sequences of Lpa and other kinase protein families from selected species using the neighbour-joining method with 1,000 bootstrap replicates. Protein homologs are indicated with an ‘H’. ob, *Oryza brachyantha*; Os-J, *Oryza sativa* L. *japonica*; Osi, *Oryza sativa* L. *indica*; Agt/Ast, *Aegilops tauschii*; sb, *Sorghum bicolor*; zm, *Zea Mays*; Si, *Setaria italica*; Ao, *Asparagus officinalis*; Ga, *Gossypium arboretum*; Cc, *Cajanus cajan*; Gm, *Glycine max*; Cm, *Cucurbita maxima*; ta, *Nicotiana tabacum*; Bn, *Brassica napus*; At, *Arabidopsis thaliana*; Ac, *Ananas cosmos*; Ztm, *Zotera marina*; Ci, *Chrysanthemum indicum*; Mf, *methanothermus fervidus*; R-570, *Saccharum*; Al, *Arabidopsis lyrata*.

Phylogenetic analysis revealed a strong relationship among the kinase proteins in the glycolysis and PA biosynthesis pathways. This suggests that Lpa proteins encoding Ins(3)P_1_ kinase are classified into the Lpa clade (Fig 6B).

### Co-segregation analysis of low PA phenotype with *lpa1-3* allele

A dCAPS assay was developed to determine the co-segregation of *lpa*1-3 allele with the low PA phenotype (Fig 7). A pair of dCAPS markers amplified a 192 bp PCR product. Digestion of this PCR product with *Taq*I yielded a 174 bp fragment in Sanggol, but an uncut fragment (192 bp) in Ilpum. Genotyping the F_2_ individuals using this dCAPS marker showed a segregation ratio, which was consistent with the expected ratio of 1:2:1 (Table 3). In addition, the dCAPS marker genotype co-segregated with the low PA phenotype in the F_2_ population. Statistical analysis using Student’s *t*-test revealed significant differences in the seed PA (S3 Fig) and P_i_ (S4 Fig) contents of Ilpum, Sanggol, and F2 individuals.

**Fig 7.**
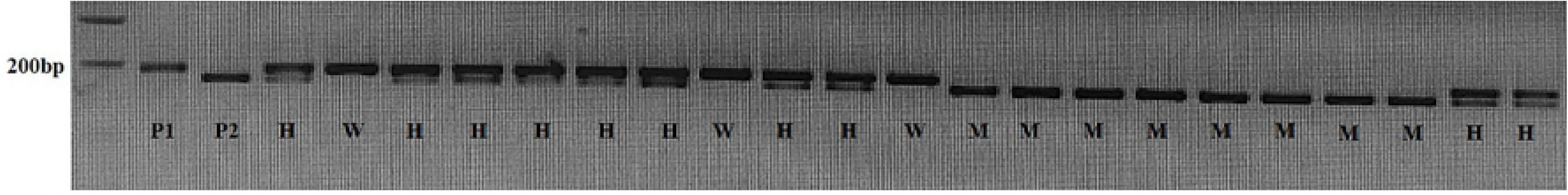
dCAPS analysis of *lp1*-3 allele in the F_2_ population derived from a cross between Sanggol and Ilpum. P1, ‘Ilpum’; P2, ‘Sanggol’; W, homozygous wild type; M, homozygous mutant; H, heterozygous.

### Biocomputational analysis

Structural analysis of Lpa1 using molecular docking of ligand and cofactors showed that Ins(3)P_1_ kinase was able to bind to the active site of Lpa1 protein, with ATP as a cofactor for catalysis (Fig 8A). Detailed view of the 3D protein model showed that Thr residue at amino acid position 208 was located in the kinase loop (Fig 8B) on the outer surface of the protein, adjacent to the entry site of the binding pocket, thus indicating its potential involvement in regulating the enzyme activity of Lpa1 protein. In addition, GPS 3.0 predicted Thr208 residue as a putative phosphorylation site, with a score of 9.66 above the cut-off value of 8.31. In previous studies, biochemical characterization of the regulatory mechanisms of various other metabolic enzymes has shown that amino acid substitutions are responsible for the reduction in enzyme activity of mutant proteins compared with wild-type proteins [47, 48]. Altogether, our results suggest that Thr208Ile amino acid substitution regulates the enzyme activity of Lpa1 protein via phosphorylation in Sanggol mutant cultivar.

**Fig 8.**
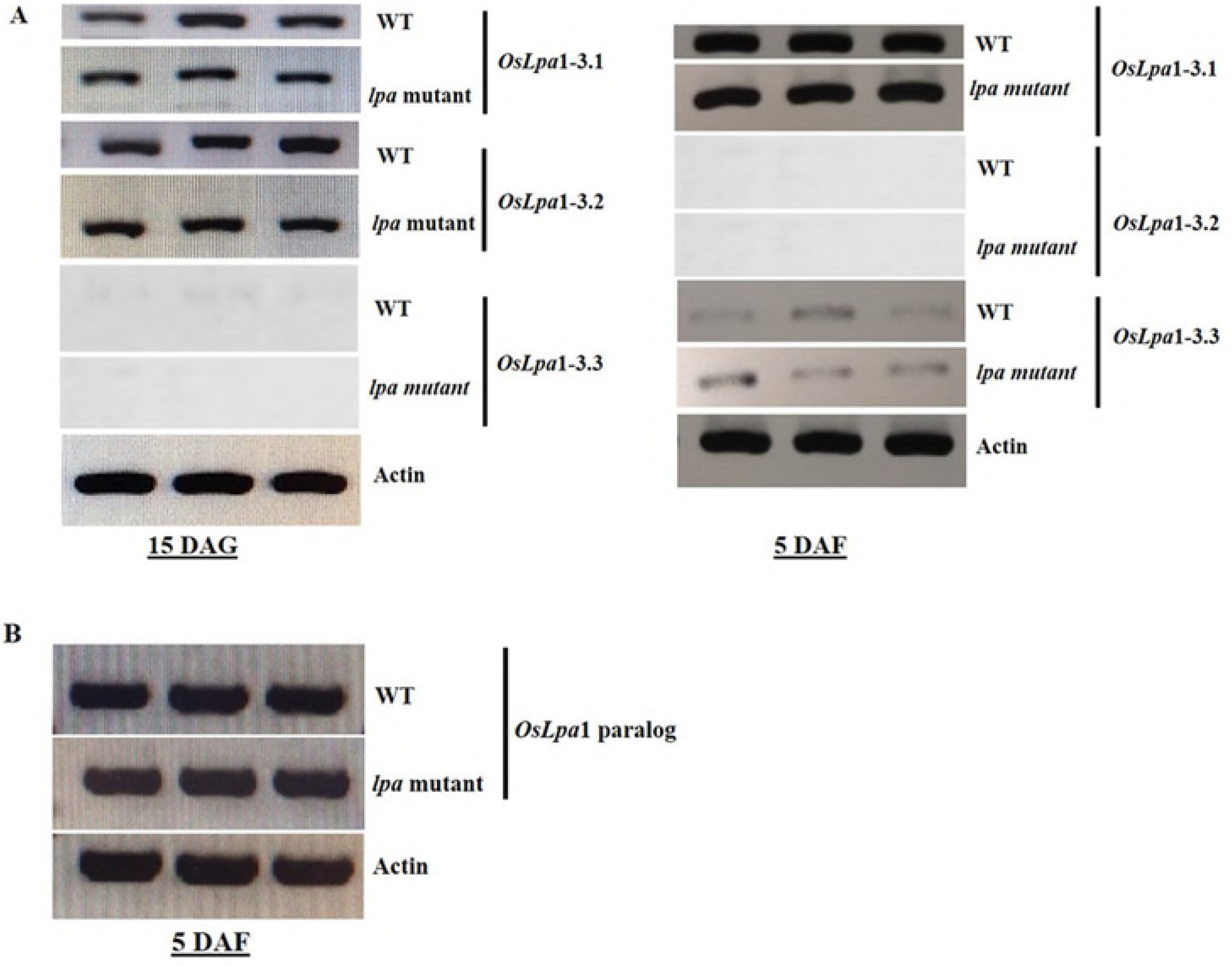
Three-dimensional (3D) model of Lpa1 protein structure. (A) 3D model of Lpa1 protein showing Ins(3)P_1_ (ligand) binding site and ATP (cofactor). (B) Thr208Ile substitution is indicated with an arrow, and the P loop containing Walker A and Walker B motifs is shown.

## Discussion

To date, several genes controlling PA biosynthesis have been reported in major crop plants [13, 20, 25, 28, 49–51]. The biosynthesis of PA proceeds via two major routes: a lipid dependent pathway, which operates in all plant tissues, and lipid independent pathway, which operates predominantly in seeds [3, 17]. In rice, molecular characterization of genes encoding MIPS, MIK, Lpa1, ITPK5/6, and IPK1 has revealed association with the low PA phenotype [21–25]. The first step of PA biosynthesis involves the conversion of D-glucose-6-phosphate to Ins(3)P_1_ by MIPS [19], which is followed by a series of phosphorylation steps, leading to the formation of InsP_6_ (S1 Fig). However, biochemical pathways leading to the conversion of Ins(3)P_1_ to InsP4, and the enzymes involved are very complex and not yet fully understood in plants [3].

Understanding the genetic basis of low PA phenotype is important for developing cultivars with low PA content in seeds. Therefore, we obtained the low PA mutant cultivar ‘Sanggol’ from Kangwon National University, Republic of Korea [31, 46]. In this study, Sanggol showed relatively poor agronomic performance compared with the wild cultivar Ilpum (Table 4). These results are in agreement with previous studies showing superior agronomic performance of wild cultivars compared with the low PA mutants [5, 14]. Edwards et al. [41] report an association between *Lpa*1 locus and grain chalkiness in rice. Similarly, the Sanggol showed high percentage of chalky grains compared with Ilpum, indicating that the low PA phenotype interacts with grain chalkiness. Thus, results of this study and previous studies suggest that the *lpa* allele plays an important role in determining the yield potential and seed quality of rice.

Phenotypic analysis using P^31^ NMR spectroscopy showed a significant reduction in PA content and an increase in P_i_ content in Sanggol seeds (Table 5). Expression analysis of PA biosynthetic genes in spikelets of the wild-type cultivar Ilpum at 5 DAF indicated that 15 genes from the lipid independent pathway were possibly responsible for the low PA content in Sanggol (Fig 3). Our data showed that a point mutation in the *OsLpa*1 locus was associated with low PA content in Sanggol seeds. Previous studies have also shown that rice low PA mutants exhibit a reduction in seed PA content because of SNPs [25, 30]. Candidate gene sequencing (Fig 4) and co-segregation analysis (Fig 7 and Table 6) confirmed that a new single recessive allele of *Lpa1*, designated as *lpa1-3*, was responsible for the low PA phenotype of Sanggol *lpa* mutant because of a C/T SNP located at nucleotide position 623 in *OsLpa*1, resulting in a single amino acid substitution (Thr208Ile). In a previous study, the *japonica* mutant ‘KBNT *lpa1-1*’ exhibited a 28% reduction in seed PA content because of a SNP (C/G to T/A), resulting in a nonsense mutation at amino acid position 409 whereas the DR1331-2 (*lpa1-2*) mutant showed a 48% reduction in seed PA content because of a single nucleotide deletion (T/A) at position 313, causing a frame shift mutation [22]. In addition, molecular characterization of the *indica* mutant ‘Os-lpa-XQZ-1’ shows the deletion of a 1,475 bp fragment in *lpa1-1*, resulting in a 38% reduction in seed PA content [30].

The *OsLpa*1 gene encodes three splice variants, all of which are expressed in seeds, suggesting that these variants play different roles in rice seed development [22, 35]. However, RT-PCR analysis of *OsLpa*1 locus revealed that *OsLpa*1-3.1 expression exhibited both vegetative and seed specificity, which indicates a major role of *OsLpa*1-3.1 in PA biosynthesis; however, *OsLpa*1-3.2 and *OsLpa*1-3.3 showed significant and dynamic changes at 15 DAG and 5 DAF, respectively (Fig 5A), suggesting that these variants play important roles in seedling growth and seed development, respectively. This finding is consistent with a previous study in rice [52]. Additionally, we investigated the expression of *OsLpa*1 paralog (Os09g0572200) and a IPK2 kinase is specific for the lipid independent pathway, *OsIpk*2 (Os02g0523800), in spikelets at 5 DAF, to provide an alternative explanation for the low level of PA in Sanggol mutant seeds. However, expression analysis suggests that the *OsLpa*1 paralog gene involved in PA biosynthesis in Sanggol (Fig 5B).

According to a previous study, *OsLpa*1 shows a weak homology to 2-PGK found in *Methanothermus fervidus* [22]. 2,3-bisphosphoglycerate (2,3-BPG), derived from 2-PGK, is a strong inhibitor of inositol polyphosphate 5-phosphatases [53]; thus, removing this inhibition may degrade inositol polyphosphate intermediates, causing a reduction in seed PA content in low PA mutants [22, 54]. Based on the structural similarity among substrates and products of *OsLpa*1 and 2-PGK, it is possible that Lpa1 protein functions as a kinase [3]. Additionally, our results revealed a single amino acid substitution (Thr208Ile) in the kinase domain of *Lpa1* in Sanggol. The *Lpa1* gene encodes Ins(3)P1 kinase, which phylogenetically groups with the Lpa clade. From the molecular docking analysis, it is evident that Ins(3)P_1_ binds to the Lpa1 protein, with ATP as a cofactor for catalysis (Fig 8A). Overall, these results suggest that Lpa1 protein functions as a kinase, and is probably involved in the conversion of Ins(3)P_1_ to *myo*-inositol 3,4-bisphosphate [Ins (3,4) P_2_].

In *Arabidopsis*, aspartic acid (Asp) to alanine (Ala) substitutions at amino acid positions 98 and 100 (Asp98Ala and Asp100Ala) in two genes encoding inositol polyphosphate kinases result in inactive enzymes and LPA phenotypes [55]. Similarly, analysis of phosphorylation deficient mutants in yeast and human shows decreased MIPS activity compared with wild- type because of amino acid substitutions at phosphorylation sites [48]. Several studies of various kinases and other metabolic enzymes show reduced enzyme activity of the mutant protein because of Thr and other amino acid substitutions at phosphorylation sites [47, 56–60]. Therefore, we speculate that a point mutation (C623T) causing Thr208Ile amino acid substitution in the loop adjacent to the entry site of the binding pocket of OsLpa1 is responsible for the altered enzyme activity of OsLpa1-3.1, resulting in reduced PA biosynthesis in Sanggol mutant seeds. Additionally, enzyme activity analysis is necessary to confirm the association of Thr208Ile substitution with the reduction in seed PA content in Sanggol low PA mutant cultivar.

Previous findings suggest that climate change and elevated CO_2_ levels negatively affect micronutrient bioavailability and total P in grains [15, 16]. However, rising CO_2_ levels are likely to increase the yield of rice crop because of the stimulation of photosynthetic rate [61]. A 1.2% increase in seed PA content under elevated CO_2_ conditions has been reported in rice [62]. In addition, crop plants need more P under elevated CO_2_ levels [63]. Soil P also exhibits a positive correlation with seed PA content in rice [64]. Limited information is available on how climate change affects seed PA content, and further studies are needed to avoid nutrient deficiencies. Reducing the seed PA content and increasing P uptake by crop plants should be a major focus of future crop breeding programs. The results of Sanggol mutant reported in this study will facilitate the development of new low PA lines with increased seed micronutrient bioavailability, high P uptake, better nutrition, and enhanced agronomic performance, despite elevated CO_2_ levels, using marker assisted introgression of the *lpa1-3* allele into elite rice varieties.

## Acknowledgments

This work was supported by a Grant from the Next-Generation BioGreen 21 Program (Plant Molecular Breeding Center number PJ013165), Rural Development Administration, Republic of Korea.

## Author Contributions

**Conceptualization:** Kishor Doddanakatte Shivaramegowda, Hee-Jong Koh

**Data curation:** Kishor Doddanakatte Shivaramegowda, Hee-Jong Koh

**Formal analysis:** Kishor Doddanakatte Shivaramegowda

**Funding acquisition:** Hee-Jong Koh

**Methodology:** Kishor Doddanakatte Shivaramegowda, Choonseok Lee, Hee-Jong Koh

**Project administration:** Hee-Jong Koh

**Resources:** Hee-Jong Koh, Soon-Kwan Hong, Jin-Kwan Ham

**Software:** Kishor Doddanakatte Shivaramegowda, Dongryung Lee, Jeonghwan Seo

**Supervision:** Hee-Jong Koh

**Validation:** Choonseok Lee

**Visualization:** Kishor Doddanakatte Shivaramegowda, Choonseok Lee

**Writing ± original draft:** Kishor Doddanakatte Shivaramegowda

**Writing, review & editing:** Kishor Doddanakatte Shivaramegowda, Choonseok Lee, Jelli Venkatesh, Zhuo Jin, Dongryung Lee, Jeonghwan Seo, Joong Hyoun Chin, Hee-Jong Koh

## Supporting information

**S1 Fig. Schematic representation of the phytic acid (PA) biosynthetic pathway.** Glu6p, glucose-6-phosphate; Ins, *myo*-inositol; PtdIns, phosphatidyl inositol; MIPS, *myo*-inositol-3-phosphate synthase; IMP, *myo*-inositol monophosphatase; MIK, *myo*-inositol kinase; LPA1, low phytic acid 1; ITPK, inositol 1,3,4-triphosphate 5/6-kinase; IPK1, inositol 1,3,4,5,6 penta*kis*phosphate 2-kinase; PIS, phosphatidyl inositol phosphate synthase; PI4K, phosphatidyl inositol 4-kinase; PIP5K, phosphatidyl inositol 4 phosphate 5-kinase; PLC, phospholipase C; IPK2, inositol 1,4,5-tris-phosphate kinase; MRP, multidrug resistance protein.

**S2 Fig. Structure of *OsLpa*1-3.2 and *OsLpa*1-3.3 splice variants.** Mutations (C53T) in the first exon of *OsLpa*1-3.2 and *OsLpa*1-3.3 are indicated with red lines. Empty boxes represent 5’ and 3’ untranslated regions (UTRs), black box represents the coding region, and lines between boxes indicate introns. ATG (start codon) and TGA (stop codon) are shown.

**S3 Fig. Statistical analysis of seed PA content among F_2_ plants derived from a cross between the mutant cultivar Sanggol and wild-type cultivar Ilpum.** Data were analyzed using the Student’s *t*-test. M, mutant; W, wild-type.

**S4 Fig. Statistical analysis of inorganic phosphorous (P_i_) content in seeds of F_2_ plants.** Data were analyzed using the Student’s *t*-test. M, mutant; W, wild-type.

**S1 Table. Primers used to sequence 16 PA biosynthetic genes in the lipid dependent and independent pathways.**

